# What would it take to describe the global diversity of parasites?

**DOI:** 10.1101/815902

**Authors:** Colin J. Carlson, Tad A. Dallas, Laura W. Alexander, Alexandra L. Phelan, Anna J. Phillips

## Abstract

How many parasites are there on Earth? Here, we use helminth parasites to highlight how little is known about parasite diversity, and how insufficient our current approach will be to describe the full scope of life on Earth. Using the largest database of host-parasite associations and one of the world’s largest parasite collections, we estimate a global total of roughly 100,000 to 350,000 species of helminth endoparasites of vertebrates, of which 85% to 95% are unknown to science. The parasites of amphibians and reptiles remain the most poorly described, but the majority of undescribed species are likely parasites of birds and bony fish. Missing species are disproportionately likely to be smaller parasites of smaller hosts in undersampled countries. At current rates, it would take centuries to comprehensively sample, collect, and name vertebrate helminths. While some have suggested that macroecology can work around existing data limitations, we argue that patterns described from a small, biased sample of diversity aren’t necessarily reliable, especially as host-parasite networks are increasingly altered by global change. In the spirit of moonshots like the Human Genome Project and the Global Virome Project, we consider the idea of a Global Parasite Project: a global effort to transform parasitology and inventory parasite diversity at an unprecedented pace.

## 1 Introduction

Parasitology is currently trapped between apparently insurmountable data limitations and the urgent need to understand how parasites will respond to global change. Parasitism is arguably the most species-rich mode of animal life on Earth (1; 2; 3), and parasites likely comprise a majority of the undescribed or undiscovered species left to modern science. (2; 4) In recent years, the global diversity and distribution of parasite richness has become a topic of particular concern (5; 6; 1), both in light of the accelerating rate of disease emergence in wildlife, livestock, and humans (7), and growing recognition of the ecological significance of many parasites. (8) Parasitic taxa are expected to face disproportionately high extinction rates in the coming century, causing a cascade of unknown but possibly massive ecological repercussions. (5; 9) Understanding the impacts of global change relies on baseline knowledge about the richness and biogeography of parasite diversity, but some groups are better studied than others. Emerging and potentially-zoonotic viruses dominate this field (10; 11; 12; 13; 14); macroparasites receive comparatively less attention.

Despite the significance of parasite biodiversity, the actual richness of most macroparasitic groups remains uncertain, due to a combination of underlying statistical challenges and universal data limitations for symbiont taxa. Particularly deserving of reassessment are helminth parasites (hereafter helminths), a polyphyletic group of parasitic worms including, but not limited to, the spiny-headed worms (acanthocephalans), tapeworms (cestodes), roundworms (nematodes), and flukes (trematodes). Helminth parasites exhibit immense diversity, tremendous ecological and epidemiological significance, and a wide host range across vertebrates, invertebrates, and plants. Estimates of helminth diversity remain controversial (1; 2; 15), especially given uncertainties arising from the small fraction of total diversity described so far (4). Though the task of describing parasite diversity has been called a “testimony to human inquisitiveness” (1), it also has practical consequences for the global task of cataloging life; one recent study proposed there could be 80 million or more species of nematode parasites of arthropods, easily reaffirming the Nematoda as a contender for the most diverse phylum on Earth. (2)

With the advent of metagenomics and bioinformatics, and the increasing digitization of natural history collections, funders are becoming interested in massive “moonshot” endeavors to catalog global diversity. Last year, the Global Virome Project was established with the stated purpose of cataloging 85% of viral diversity within vertebrates (particularly mammals and birds, which host almost all emerging zoonoses), with an investment of $1.2 billion over 10 years. Whereas the Global Virome Project is ultimately an endeavor to prevent the future emergence of the highest-risk potential zoonoses—the natural evolution of decades of pandemic-oriented work at the edge of ecology, virology, and epidemiology—we suggest parasitologists have the opportunity to set a more inclusive goal. Between a quarter and half of named virus species can infect humans (14), while human helminthiases are a small, almost negligible fraction of total parasite diversity despite their massive global health burden. The need to understand global parasite diversity reflects a more basic set of questions about the world we live in, and the breadth of life within it.

Here we ask, what it would take to completely describe global helminth diversity in vertebrates? The answer is just as dependent on how many helminth species exist as it is on the rate and efficiency of parasite taxonomic description efforts. We set out to address three questions:

I. What do we know about the global process of describing and documenting parasite biodiversity, and how will it change in the future?
II. How many helminth species should we expect globally, and how much of that diversity is described?
III. How many years are we from describing all of global parasite diversity, and what can (and can’t) we do with what we have?

From there, we make recommendations about where the next decade of parasite systematics and ecology might take us.

## 2 The data

To answer all three questions, we take advantage of two collections-based datasets that have been made available in the last decade (Figure 1). The biological collections housed at museums, academic research institutions, and various private locations around the world are one of the most significant “big data” sources for biodiversity research (16), especially for parasites. (17; 18) The Natural History Museum in London, UK (NHM) curates the Host-Parasite Database, which includes regional lists of helminth-host associations, including full taxonomic citations for helminth species. (19; 20) By species counts alone, the NHM dataset is perhaps the largest species interaction dataset published so far in ecological literature. (6) In our updated scrape of the web interface, which will be the most detailed version of the dataset ever made public, there are a raw total of 109,060 associations recorded between 25,740 helminth species (including monogeneans) and 19,097 hosts (vertebrate and invertebrate).

**Figure 1:**
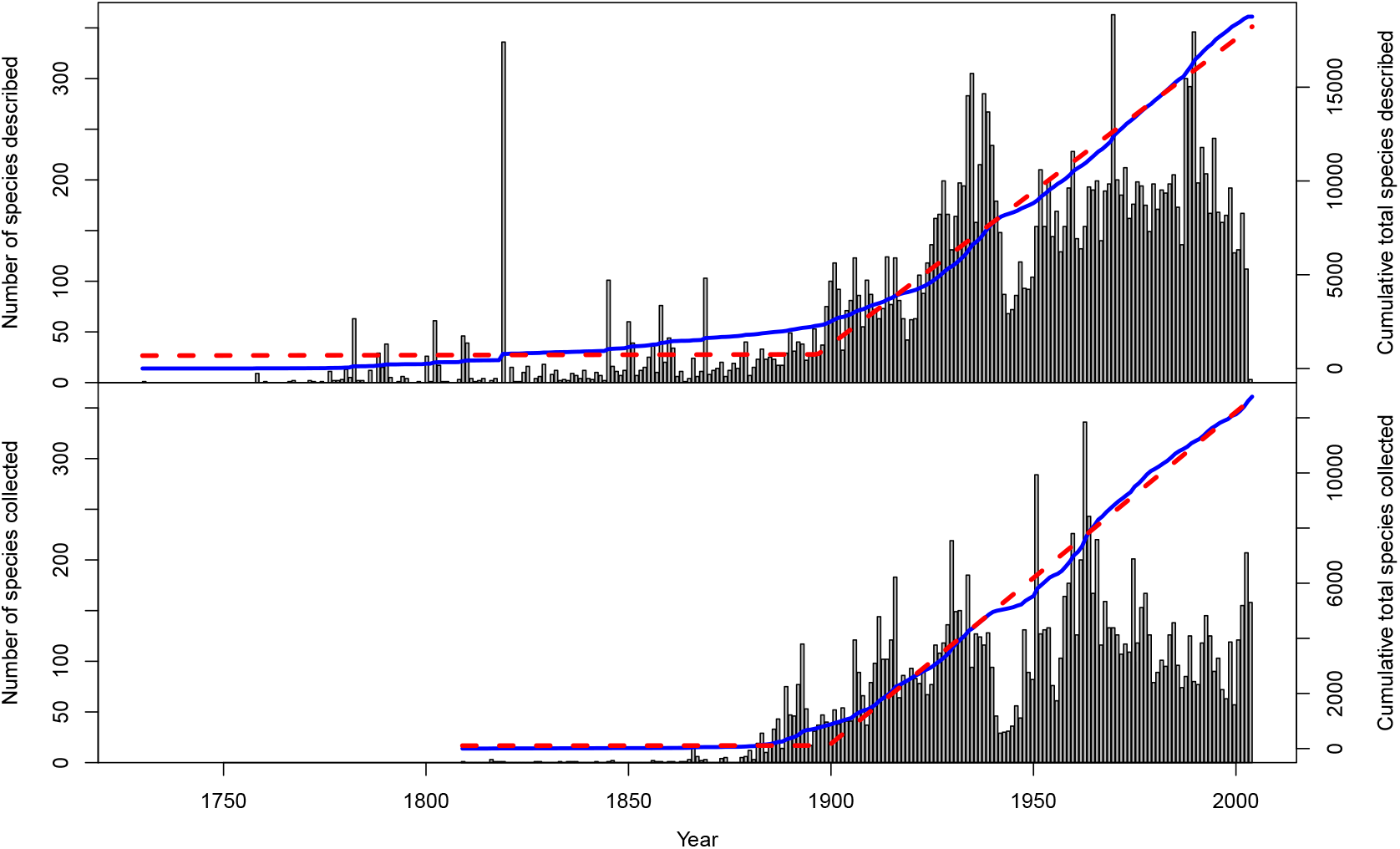
Rates of helminth descriptions (top, from NHM data) and collections (bottom, from the USNPC). Blue trends indicate cumulative totals, and red lines give a breakpoint regression with a single breakpoint (1912 for the NHM data, 1903 for the USNPC data). Although the current trend appears to be leveling off, it is unlikely this indicates a saturating process (as comparably illustrated by the drop in sampling during the Second World War, 1940-1945).

The U.S. National Parasite Collection (USNPC) is one of the largest parasite collections in the world, and is one of the most significant resources used by systematists to discover, describe, and document new species (17; 21). The published records constitute the largest open museum collection database for helminths, especially in terms of georeferenced data availability (5). Here, we use a recent copy of the USNPC database that includes 89,580 specimen records, including 13,426 species recorded in the groups Acanthocephala, Nematoda, and Platyhelminthes. (Of these we assume the vast majority are vertebrate parasites.) In combination, the two datasets represent the growing availability of big data in parasitology, and allow us to characterize parasite diversity much more precisely than we could have a decade ago.

## 3 How does parasite biodiversity data accumulate?

Describing the global diversity of parasites involves two major processes: documenting and describing diversity through species descriptions, geographic distributions, host associations, etc.; and consolidating and digitizing lists of valid taxonomic names and synonyms (e.g. ITIS, Catalogue of Life, WoRMS). Both efforts are important, time-consuming, and appear especially difficult for parasites.

### 3.1 Why has helminth diversity been so difficult to catalogue?

The most obvious reason is the hyperdiversity of groups like the Nematoda, but this only tells part of the story. Other hyperdiverse groups, like the sunflower family (Asteraceae), have far more certain richness estimates (and higher description rates) despite being comparably speciose. Several hypotheses are plausible: surveys could be poorly optimized for the geographic and phylogenetic distribution of helminth richness, or remaining species might be objectively harder to discover and describe than known ones were. Perhaps the most popular explanation is that taxonomists’ and systematists’ availability might be the limiting factor (22; 23); the process of describing helminth diversity relies on the dedicated work of systematic biologists, and the availability and maintenance of long-term natural history collections. However, Costello *et al*. (24) observed that the number of systematists describing parasites has increased steadily since the 1960s, with apparently diminishing returns. Costello posited this was evidence the effort to describe parasites has reached the “inflection point,” with more than half of all parasites described; this assessment disagrees noticeably with many others in the literature. (23)

### 3.2 Have we actually passed the inflection point?

No, probably not. We show this by building species accumulation curves over time, from two different sources: the dates given in taxonomic authority citations in the NHM data, and the date of first accession in the USNPC data, for each species in the dataset (Figure 1). Both are a representation of total taxonomic effort, and vary substantially between years. Some historical influences are obvious, such as a drop during World War II (1939-1945). Recently the number of parasites accessioned has dropped slightly, but it seems unlikely (especially given historical parallels) that this reflects a real inflection point in parasite sampling, and is probably instead reflects a limitation of the data structure; the NHM data in particular has not been updated since 2013. Despite interannual variation, the accumulation curves both demonstrate a clear cut pattern: sometime around the turn of the 20th century, they turn upward and increase linearly. Since 1897, an average of 163 helminth species have been described annually (*R*^2^ = 0.991, *p* < 0.001), while an average of 120 species are added to collections every year since 1899 (*R*^2^ = 0.998, p < 0.001). The lack of slowing down in those linear trends is a strong indicator that we remain a long way from a complete catalog of helminth diversity.

### 3.3 Are we looking in the wrong places?

An alternate explanation for the slow rate of parasite discovery is that the majority of parasite diversity is in countries where sampling effort is lower, and *vice versa* most sampling effort and research institutions are in places with more described parasite fauna (25). Recent evidence suggests species discovery efforts so far have been poorly optimized for the underlying—but mostly hypothetical—richness patterns of different helminth groups. (25; 26) Ecologists have started to ask questions that could help optimize sampling: do parasites follow the conventional latitudinal diversity gradient? Are there unique hotspots of parasite diversity, or does parasite diversity peak in host biodiversity hotspots? (27; 6; 1; 25; 28; 29) But our ability to answer these types of questions is predicated on our confidence that observed macroecological patterns in a small (and uncertain) percentage of the world’s helminths are representative of the whole.

### 3.4 Are species described later qualitatively different?

If helminth descriptions have been significantly biased by species’ ecology, this should produce quantitative differences between the species that have and haven’t yet been described. We examine two easily intuited sources of bias: body size (larger hosts and parasites are better sampled) and host specificity (generalist parasites should be detected and described sooner). We found a small but highly significant trend of decreasing body size for both hosts and parasites, suggesting the existence of a sampling bias, but not necessarily suggesting unsampled species should be massively different. (Figure 2) For host specificity, we find an obvious pattern relative to description rates, though less so for collections data. (Figure 3) The inflection point around 1840 is likely a byproduct of the history of taxonomy, as the *Series of Propositions for Rendering the Nomenclature of Zoology Uniform and Permanent*—now the International Code for Zoological Nomenclature—was first proposed in 1842, leading to a standardization of host nomenclature and consolidation of the proliferation of multiple names for single species.

**Figure 2:**
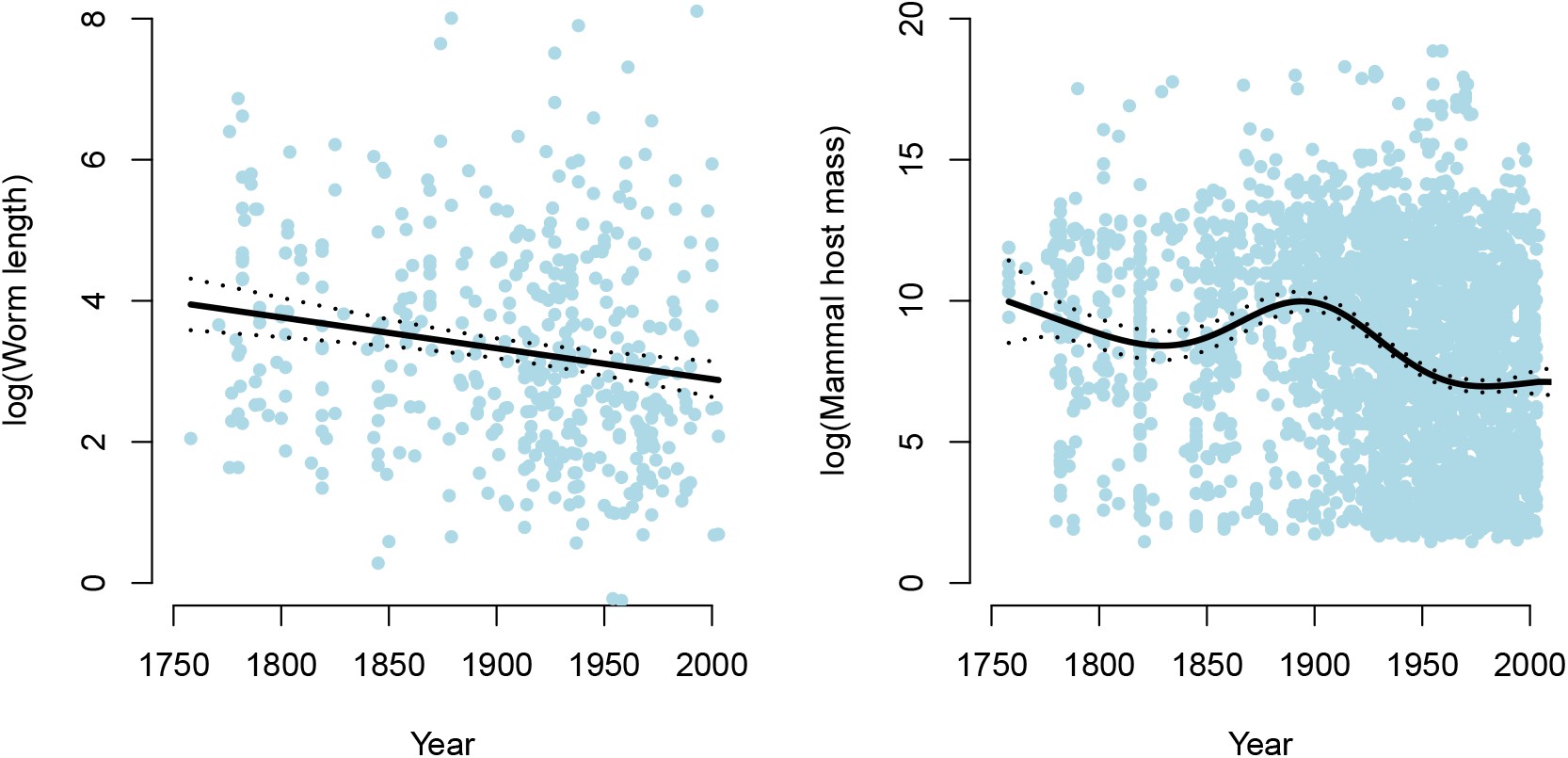
We found evidence of weak but highly significant declines over time in parasite adult body length (left; smooth term p = 0.0003) and host body size across known host associations (right; smooth term p < 0.0001). This confirms a mild description bias for larger parasites in larger hosts.

**Figure 3:**
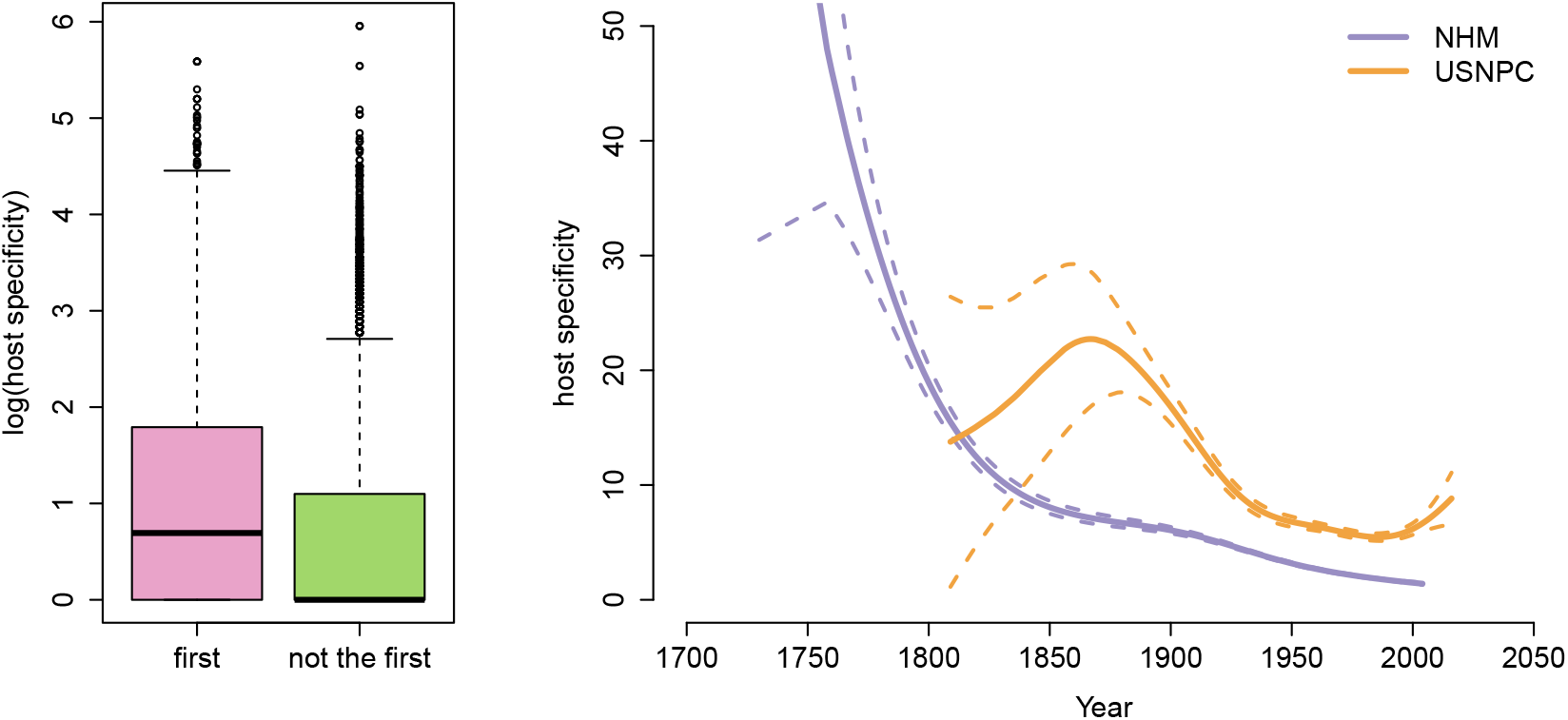
The type species (the first described in a genus) has a statistically significantly higher average host specificity than those that follow. Parasites described earlier typically have a higher degree of generalism, especially prior to the 1840s; specimens collected after roughly the 1870s also apparently tend towards more host-specific species than those from older collections. (Curves are generalized additive models fit assuming a negative binominal distribution, with dashed lines for the 95% confidence bounds.)

The temporal trend also likely reflects the history of taxonomic revisions, as the first species reported in a genus tends to have a higher range of hosts, morphology, and geography, while subsequent revisions parse these out into more appropriate, narrower descriptions. Using the NHM data, we can easily show that the first species reported in every genus (usually the type species but not always, given incomplete sampling) generally has significantly higher reported numbers of hosts (Wilcoxon rank sum test: *W* = 22, 390, 629, *p* < 0.001; Figure 3). This is because type species often become umbrella descriptors that are subsequently split into more species after further investigation, each with only a subset of the initial total host range. Based on our results, we can expect undescribed species of helminths to be disproportionately host-specific.

## 4 How many helminths?

### 4.1 How do we count parasites?

For many groups of parasites, the number of species known to science is still growing exponentially, preventing estimation based on the asymptote of sampling curves. (30) In some cases, there are workarounds: for example, the diversity of parasitoid wasps (Hymenoptera: Braconidae) has been estimated based on the distribution of taxonomic revisions rather than descriptions. (31) But for helminths, every major estimate of diversity is based on the scaling between host and parasite richness, a near-universal pattern across spatial scales and taxonomic groups. (32; 33; 6) The scaling of hosts and fully host-specific parasites can be assumed to be linear: for example, every arthropod is estimated to have at least one host-specific nematode. (2) Poulin and Morand (30) proposed an intuitive correction for generalists:

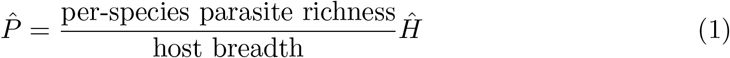

Poulin and Morand (30) compiled independently-sourced estimates of host specificity and per-species richness, and the resulting estimate of ~75,000 to 300,000 helminth species was canon for a decade. (1) However, Strona and Fattorini (15) used the NHM data to show that subsampling a host-parasite network approximately generates power law scaling, not linear scaling, which reduced estimates by of helminth diversity (in helminth and vertebrate taxon pairs) by an average of 58%. However, they made no overall corrected estimate of helminth diversity in vertebrates.

### 4.2 What do we know now that we didn’t before?

Examining bipartite host-affiliate networks across several types of symbiosis, including the vertebrate-helminth network (from the NHM data), we previously found approximate power law behavior in every scaling curve. (14) The underlying reasons for this pattern are difficult to ascertain, and may or may not be connected to approximate power-law degree distributions in the networks. Regardless, the method seems to work as a tool for estimating richness; using the new `R` package `codependent` (34), we used these tools to show that viral diversity in mammals is probably only about 2-3% of the estimates generated with linear extrapolation by the Global Virome Project. (14) Here, we build on this work by showing how association data can be used to estimate the proportion of overlap among groups, and thereby correct when adding together parasite richness sub-totals. (See **Materials and Methods.**)

### 4.3 How many species are there?

Building on previous studies (1; 15), we re-estimated global helminth diversity using `codependent`, a taxonomically-updated version of the NHM dataset, and a new formula for combining parasite richness across groups. (Table 1) In total, we estimated 103,078 species of helminth parasites of vertebrates, most strongly represented by trematodes (44, 262), followed by nematodes (28,844), cestodes (23,749), and acanthocephalans (6,223). Using an updated estimate of bony fish richness significantly increased these estimates from previous ones, with over 37,000 helminth species in this clade alone. Birds and fish were estimated to harbor the most helminth richness, but reptiles and amphibians had the highest proportion of undescribed diversity. The best-described groups were nematode parasites of mammals (possibly because so many are zoonotic and livestock diseases) and cestode parasites of the cartilaginous fishes (perhaps due to the expertise of a strong collaborative research community, including the participants in the Planetary Biodiversity Inventory project on cestode systematics). (35)

**Table 1:** Helminth diversity, re-estimated: How many helminth species (top), and what percentage of species have been described (bottom)?

|  | Chondrichthyes | Osteichthyes | Amphibia | Reptilia | Aves | Mammalia | Total |
| --- | --- | --- | --- | --- | --- | --- | --- |
| Acanthocephala | 169<br>(4%) | 3,572<br>(13%) | 765<br>(3%) | 785<br>(4%) | 1,184<br>(14%) | 886<br>(12%) | 6,223<br>(11%) |
| Cestoda | 2,108<br>(28%) | 5,875<br>(12%) | 637<br>(5%) | 2,153<br>(5%) | 10,257<br>(14%) | 4,061<br>(26%) | 23,749<br>(16%) |
| Nematoda | 566<br>(14%) | 10,712<br>(11%) | 2,148<br>(10%) | 4,537<br>(12%) | 3,925<br>(19%) | 7,902<br>(30%) | 28,844<br>(17%) |
| Trematoda | 391<br>(16%) | 17,745<br>(19%) | 3,700<br>(6%) | 12,153<br>(4%) | 8,778<br>(17%) | 4,550<br>(23%) | 44,262<br>(14%) |
| Total | 3,234<br>(23%) | 37,904<br>(15%) | 7,250<br>(7%) | 19,628<br>(6%) | 24,144<br>(16%) | 17,399<br>(26%) | 103,078<br>(15%) |

### 4.4 Do we trust these estimates?

Although estimates from a decade ago were surprisingly close given methodological differences (1), we now have a much greater degree of confidence in our overall estimate of vertebrate helminth richness. However, some points of remaining bias are immediately obvious. The largest is methodological: by fitting power law curves over host richness, we assumed all hosts had at least one parasite from any given helminth group. While this assumption worked well for mammal viruses, it may be more suspect especially for the less-speciose groups like Acanthocephala. On the other hand, the power law method is prone to overestimation in several ways enumerated in (14). Furthermore, Dallas *et al*. (36) estimated that 20-40% of the host range of parasites is underdocumented in the Global Mammal Parasite Database, a sparser but comparable dataset. If these links were recorded in our data, they would substantially expand the level of host-sharing and cause a reduction of the scaling exponent of power laws, causing lower estimates. On the other hand, if we know that the majority of undescribed parasite diversity is far more host specific than known species, our estimates would severely underestimate in this regard. At present, it is essentially impossible to estimate the sign of the these errors once compounded together.

### 4.5 What about cryptic diversity?

One major outstanding problem is cryptic diversity, the fraction of undescribed species that are genetically distinct but morphologically indistinguishable, or at least so subtly different that their description poses a challenge. Many of the undescribed species could fall in this category, and splitting them out might decrease the apparent host range of most species, further increasing estimates of total diversity. Dobson *et al*. (1) addressed this problem by assuming that the true diversity of helminths might be double and double again their estimate; while this makes sense conceptually, it lacks any data-driven support. The diversity of cryptic species is unlikely to be distributed equally among all groups; for example, long-standing evidence suggests it may be disproportionately higher for trematodes than cestodes or nematodes. (37)

We can loosely correct our overall richness estimates for cryptic diversity. A recently-compiled meta-analysis suggests an average of 2.6 cryptic species per species of acanthocephalan, 2.4 per species of cestode, 1.2 per species of nematode, and 3.1 per species of digenean. (38) Using these numbers, we could push our total estimates to at most 22,404 acanthocephalan species, 80,747 cestodes, 63,457 nematodes, and a whopping 181,474 species of trematodes, with a total of 348,082 species of helminths. However, there may be publication bias that favors higher cryptic species rates (or at least, zeros may be artificially rare), making these likely overestimates. Increased sampling will push estimates higher for many species, and eventually will allow a more statistically certain estimate of the cryptic species “multiplication factor” needed to update the estimates we present here.

## 5 Could we describe the world’s parasite diversity?

### 5.1 How long would it take to catalog global helminth diversity?

We estimated 103,079 total helminth species on Earth, of which 13,426 (13.0%) are in the USNPC and 15,817 (15.3%) are in the NHM Database. At the current rates we estimated, it would take 536 years to describe global helminth diversity and catalog at least some host associations (based on the NHM data as a taxonomic reference), and 745 years to add every species to the collection (based on the USNPC). Including the full range of possible cryptic species would push the total richness to 348,082 helminth species (95% undescribed), which would require 2,040 years to describe and 2,779 years to collect.

Even with hypothetical overcorrections, these are daunting numbers: for example, if the NHM only captures one tenth of known helminth diversity, and thereby underestimates the rate of description by an order of magnitude, it would still take two centuries to describe remaining diversity. These estimates are also conservative in several ways: the majority of remaining species will be more host-specific and therefore harder to discover, and the process would almost certainly undergo an asymptote or at least a mild saturating process. Moreover, many of the 13,426 unique identifiers in the USNPC are either currently or may be synonyms of valid names and may be corrected through taxonomic revision and redetermination; previous estimates suggest invalid names may outnumber valid ones, in some data. (24)

### 5.2 Where is the undescribed diversity?

Previous work has argued that current patterns of helminth description are poorly matched to underlying richness patterns, though those patterns are also unknown and assumed to broadly correspond to host biodiversity (25). Here, we used the scaling between host and parasite diversity to predict the “maximum possible” number of parasites expected for a country’s mammal fauna, and compared that to known helminths described from mammals in the NHM dataset (Figure 4). While these estimates are liberal in the sense that they include the global range of parasite fauna associated with given hosts, they are also conservative in that they are uncorrected for cryptic diversity, or the possibility of higher host specificity in the tropics.

**Figure 4:**
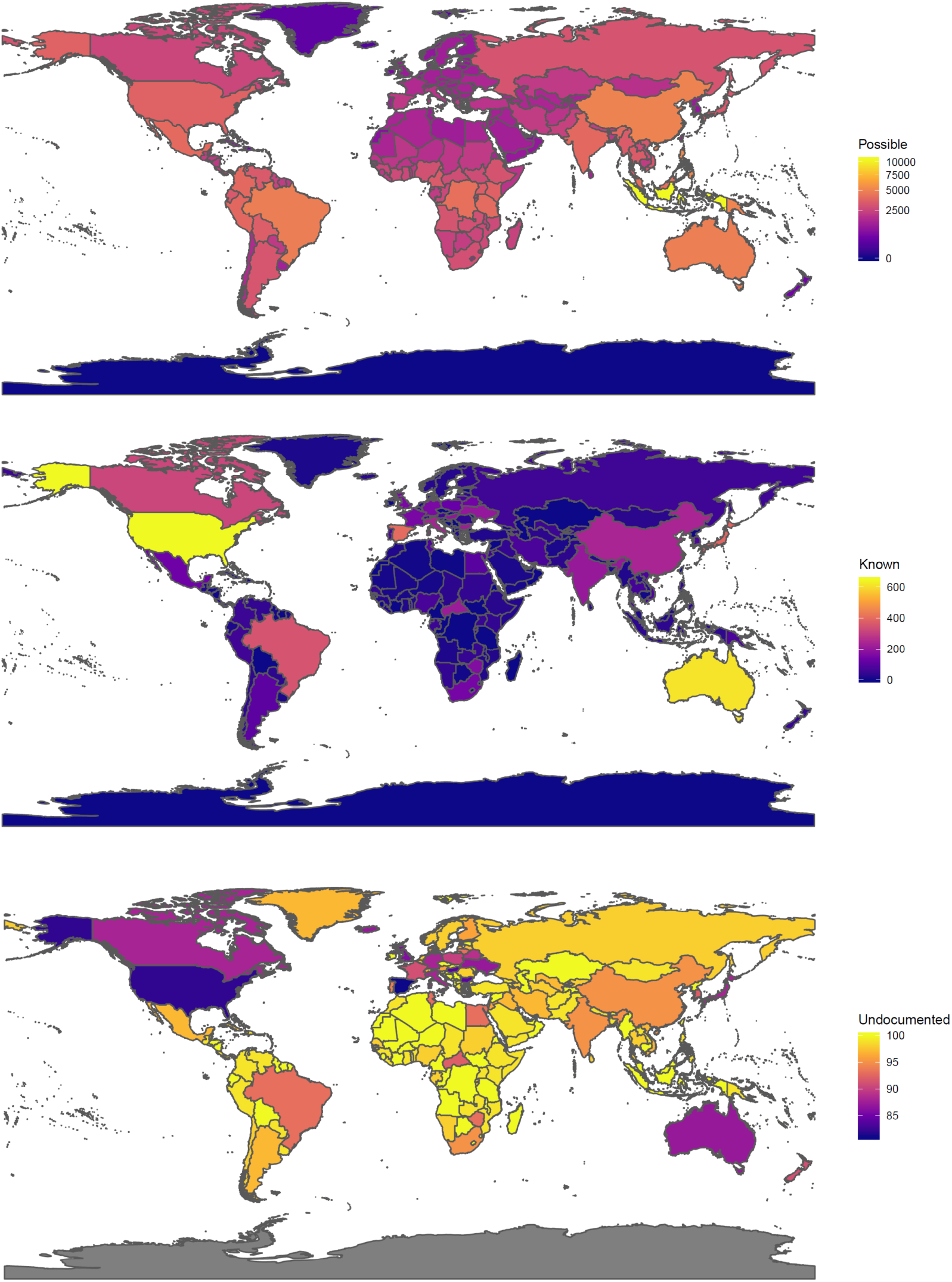
The distribution of maximum possible helminth richness in mammals (top), the number of known helminth parasites of mammals as recorded by country in the NHM data (middle), and the maximum percentage of undocumented helminth fauna by country (bottom).

We found that helminths were best known in the handful of countries that dominate parasite systematics work (the United States, Australia, Brazil, Canada, China, and some European countries). But even in these places, most species are probably undescribed; many countries have no records at all, including large countries like the Democratic Republic of the Congo that are mammal diversity hotspots. Between 80% and 100% of possible parasite diversity could be locally undescribed for most of the world—high estimates, but plausible given a global undescribed rate of 85-95%. This spatial pattern likely reflects a combination of language and access barriers (data in Chinese and Russian collections, for example, are known to be substantial, but inaccessible to our present work), and a broader inequity arising from the concentration of institutions and researchers in wealthy countries, and the corresponding disproportionate geographic focus of research. (39) Previous research has noted that African parasitology has been especially dominated by foreign researchers (40), and African parasitologists remain particularly underrepresented in Western research societies. (41)

### 5.3 How much can we do with what we have?

Or, to put the question another way: With such a small fraction of parasite diversity described, how confident can we be in macroecological patterns? A parallel problem was encountered by Quicke (42) as part of a longer-term effort to estimate global parasitoid wasp diversity. (43; 31) Only a year after publishing a paper (44) exploring similar macroecological patterns to those we have previously explored (6; 45), Quicke concluded “we know too little” to make conclusions about macroecological patterns like latitudinal trends. (42) For parasitoid wasps, the problem is attributable to a similar set of systemic biases, like underdescription of tropical fauna, or a bias in species description rates towards larger species first.

Given that almost 90% of helminth diversity is undescribed (and closer to 100% is undescribed in many places), parasite ecologists need to approach work with “big data” with a similar degree of caution. Working at the level of ecosystems or narrowly-defined taxonomic groups may help sidestep some of these issues.(28) But at the global level, patterns like a latitudinal diversity gradient could be the consequence of real underlying trends, or just as easily be the consequence of extreme spatial sampling bias in collections and taxonomic descriptions and revisions.

It will take decades or even centuries before datasets improve substantially enough to change our degree of confidence in existing macroecological hypotheses. Given this problem, Poulin (23) recommended abandoning the task of estimating parasite diversity, and assuming parasite richness is determined “simply [by] local host species richness.” However, at global scales, this is not necessarily supported (46); Dallas *et al*. (6) showed that the per-host richness of parasite fauna varied over an order of magnitude across different countries in the NHM data, a spatial pattern with little correlation to mammal biodiversity gradients. Even this result is nearly impossible to disentangle from sampling incompleteness and sampling bias. Moreover, even at mesoscales where “host diversity begets parasite diversity” is usually a reliable pattern, anthropogenic impacts are already starting to decouple these patterns (47). At the present moment, helminth richness patterns could be functionally unknowable at the global scale. The same is likely true of many other groups of metazoan parasites that are far more poorly described.

## 6 The case for a Global Parasite Project

Given the extensive diversity of helminths, some researchers have argued in favor of abandoning the goal of ever fully measuring *or* cataloging parasite diversity, focusing instead on more “practical” problems. (23) At current rates of description, this is a reasonable outlook; even with several sources of unquantifiable error built into our estimates, it might seem impossible to make a dent within a generation. However, we dispute the idea that nothing can be done to accelerate parasite discovery. Funding and support for most scientific endeavors are at an unprecedented high in the 21^st^ Century. Other scientific moonshots, from the Human Genome Project to the Event Horizon Telescope image of the M87 black hole, would have seemed impossible within living memory.

For parasitology, the nature and urgency of the problem call for a similarly unprecedented effort. For some purposes, the 5–15% of diversity described may be adequate to form and test ecoevolutionary hypotheses. But the reliability and accuracy of these data will become more uncertain in the face of global change, which will re-assemble hostparasite interactions on a scale that is nearly impossible to predict today. As climate change progresses, an increasing amount of our time and energy will be spent attempting to differentiate ecological signals from noise *and* anthropogenic signals. Though some consider the task of cataloging parasite diversity a “testimony to human inquisitiveness” (1), it is also a critical baseline for understanding biological interactions in a world on the brink of ecological collapse. Along the same lines of the Global Virome Project, we suggest that parasitology is ready for a “Global Parasite Project”: an internationally-coordinated effort to revolutionize the process of cataloging parasite diversity.

Although many parasitic clades would be worth including in a Global Parasite Project, helminths provide an invaluable model for several key points. First, modern methods make it possible to set realistic and tangible targets, and budget accordingly. Recently, the global parasite conservation plan (48) proposed an ambitious goal of describing 50% of parasite diversity in the next decade. From the bipartite rarefaction method (15; 14), we can back-estimate how many hosts we expect to randomly sample before we reach that target. For example, describing 50% of terrestrial nematode parasites would require sampling 3,215 new reptile host species, 2,560 birds, 2,325 amphibians, and only 995 mammals. These estimates assume diversity accumulates randomly, and hosts are sampled in an uninformed way. In practice, with knowledge about existing ecological and geographic biases, we can target sampling to accelerate species discovery, just as previous programs like the Planetary Biodiversity Inventory tapeworm project have, to great success. (35)

Second, any moonshot effort to describe parasite diversity would have to start with museums and collections. Systematics is the backbone of biodiversity science (49; 50), and especially in parasitology, collections are the backbone of systematics. (22; 51) They are also some of the most vulnerable research institutions in modern science: collections are chronically underfunded and understaffed, sometimes to the point of dissolving. Even well-funded collections are still mostly undigitized, ungeoreferenced, and unsequenced (17), and massive volumes of “grey data” are unaccounted for in collections that are isolated from the global research community, or fall on opposite sides of deep historical divides (e.g., between Soviet and American science). In all likelihood, hundreds or thousands of parasite species have already been identified and are waiting to be described from museum backlogs, or their descriptions have been recorded in sources inaccessible due to digital access, language barriers, and paywalls. Technological advances in the coming decade—like faster bioinformatic pipelines for digitization, easier DNA extraction from formalin-fixed samples, or cryostorage of genomic-grade samples—will expand the possibilities of collections-based work, but are insufficient to fix many of the structural problems in the field.

Whereas the proposed Global Virome Project has focused mostly on capacity building for field sampling and labwork, a Global Parasite Project could probably achieve comparable rates of parasite description (on a lower budget) by focusing on collections science. If the existing research and funding model continues into the next decade, most “available” parasite data will be collected by Western scientists running field trips or long-term ecological monitoring programs that mostly feed into collections at their home institutions. Building out American and European parasite collections with globally-sourced specimens would only perpetuate existing data gaps and research inefficiencies, and the structural inequities and injustices they reflect. Increasingly, biomedical research is under legitimate scrutiny for parachute research—Western-driven research “partnerships” that leverage international project design for exploitative and extractive sampling, with little benefit to partners in the Global South (52; 53; 54). Though our hypothetical Global Parasite Project would be focused primarily on ecology, rather than biomedical or global health priorities, systematics and conservation are no exception to these conversations.

A Global Parasite Project, and its governance principles, would need to focus on supporting collections work and strengthening infrastructure around the world, with explicit priority on equity and local leadership. Recent developments in international law are particularly relevant to this end. (55) The Nagoya Protocol on Access to Genetic Resources and the Fair and Equitable Sharing of Benefits Arising from their Utilization to the Convention on Biological Diversity (Nagoya Protocol) establishes a regime to ensure that access to genetic resources—which some countries may define to include parasites— is coupled with the equitable sharing of benefits from their use. While implementation of the Nagoya Protocol varies between countries, it codifies important norms addressing injustices in obtaining parasites for collections, and inequities in the benefits arising directly or indirectly from their use, which may include capacity building, technology transfer, and recognition in scientific publications.

Done right, a Global Parasite Project would build resilient capacities for local priorities, through financial and technical support that empowers local researchers in resource-constrained settings. The support provided could include a combination of training, funding, conferences and meetings, and technology transfer. These can be identified on a case-by-case basis to meet local priorities, which could include formalizing parasite collections, in cases where the component collections are distributed across departments; improving or modernizing specimen preservation methods or physical infrastructure; and digitizing and sequencing collections. (35; 56) Following these steps could fill major data gaps, and make collections around the world more resistant to damage, disasters, and gaps in research support. In turn, there is a wealth of local technical knowledge and expertise in countries where parasite collections are underserved. This is an opportunity for locally-led, multilateral capacity-building, and, where appropriate, dissemination of local knowledge to the broader scientific community with clear principles for locally-led publications and clear attribution. This work should expand avenues for parasitologists in the Global South to be recognized and engaged as active participants in the global research community.

Third, a Global Parasite Project would need to focus not just on completeness in parasite descriptions, but in host-parasite interaction data. The sparseness of existing network datasets can make estimates of affiliate diversity an order of magnitude more uncertain (14), and describing new parasites as fast as possible might make this problem more pronounced. An active effort needs to be made to fill in the 20-40% of missing links in association matrices, potentially using model-predicted links to optimize sampling (36). Better characterizing the full host-parasite network would have major benefits for actionable science, ranging from the triage process for parasite conservation assessments (48), to work exploring the apparently-emerging sylvatic niche of Guinea worm and its implications for disease eradication (57).

This is where ecologists fit best into a parasite moonshot. Rather than establishing an entirely novel global infrastructure for field research, we can fund a major expansion of parasitology in existing biodiversity inventories. The vast majority of animals already collected by field biologists have easily-documented symbionts, which are nevertheless neglected or discarded during sampling. In response, recent work has suggested widespread adoption of integrative protocols for how to collect and document the entire symbiont fauna of animal specimens. (58; 59) Building these protocols into more biodiversity inventories will help capture several groups of arthropod, helminth, protozoan, and fungal parasites, without unique or redundant sampling programs for each. In cases where destructive sampling is challenging (rare or elusive species) or prohibitive (endangered or protected species), nanopore sequencing and metagenomics may increasingly be used to fill sampling gaps. Collecting data these ways will improve detection of parasites’ full host range, and allow researchers to explore emerging questions about how parasite metacommunities form and interact. (60) As novel biotic interactions form and are detected in real-time, this could become a major building block of global change research. (48)

Despite decades of work calling out the shortage of parasitologists and the “death” of systematics (61; 22), the vast diversity of undescribed parasites has never stopped the thousands of taxonomists and systematists who compiled our datasets over the last century—mostly without access to modern luxuries like digital collections or nanopore sequencing. A testimony to persistence and resourcefulness, these data provide the roadmap for a new transformative effort to describe life on Earth. In an era of massive scientific endeavours, a coordinated effort to describe the world’s parasite diversity seems more possible than ever. There may never be a Global Parasite Project *per se*, but the current moment may be the closest we’ve ever been to the “right time” to try for one. If biologists want to understand how the entire biosphere is responding to a period of unprecedented change, there is simply no alternative.

## Acknowledgements

Thanks to Shweta Bansal, Phil Staniczenko, and Joy Vaz for formative conversations, and to the Georgetown Environment Initiative for funding.

## Materials and Methods

### Data Assembly and Cleaning

The data we use in this study comes from two sources: the U.S. National Parasite Collection, and the London Natural History Museum’s host-parasite database. We describe the cleaning process for both of these sources in turn. All data, and all code, are available on Github at github.com/cjcarlson/helminths.

The U.S. National Parasite Collection has been housed at the Smithsonian National Museum of Natural History since 2013, and is one of the largest parasite collections in the world. The collection is largely digitized and has previously been used for global ecological studies. (5) We downloaded the collections database from EMu in September 2017. The collection includes several major parasitic groups, not just helminths, and so we filtered data down to Acanthocephala, Nematoda, and Platyhelminthes. Meta-data associated with the collection has variable quality, and host information is mostly unstandardized, so we minimize its use here.

The London Natural History Museum’s host-parasite database is an association list for helminths and their host associations, dating back to the Host-Parasite Catalogue compiled by H.A. Baylis starting in 1922. The database itself is around 250,000 unique, mostly location-specific association records digitzed from a reported 28,000 scientific studies. The NHM dataset has been used for ecological analysis in previous publications (6; 62; 63), but here we used an updated scrape of the online interface to the database. Whereas previous work has scraped association data by locality, we scraped by parasite species list from previous scrapes, allowing records without locality data to be included, and therefore including a more complete sample of hosts. The total raw dataset comprised 100,370 host-parasite associations (no duplication by locality or other metadata), including 17,725 hosts and 21,115 parasites.

We cleaned the NHM data with a handful of validation steps. First, we removed all host and parasite species with no epithet (recorded as “sp.”), and removed all pre-revision name parentheticals. We then ran host taxonomy through ITIS with the help of the `taxize` package in R, and updated names where possible. This also allowed us to manually re-classify host names by taxonomic grouping. Parasite names were not validated because most parasitic groups are severely under-represented (or outdated) in taxonomic repositories like WORMS and ITIS. At present, no universal, reliable dataset exists for validating parasite taxonomy. After cleaning, there were a total of 13,162 host species and 20,016 parasite species with a total of 73,273 unique interactions; this is compared to, in older scrapes, what would have been a processed total of 61,397 interactions among 18,583 parasites and 11,749 hosts. We finally validated all terrestrial localities by updating to ISO3 standard, including island territories of countries like the United Kingdom; many localities stored in the NHM data predate the fall of the USSR or are have similar anachronisms.

### Trends over Time

#### Description rates

In the NHM data, we assigned dates of description by extracting year from the full taxonomic record of any given species (e.g., *Ascaris lumbricoides* Linnaeus, 1758) using regular expressions; in the USNPC data, we extracted year from the accession date recorded for a given specimen. We added together the total number of species described (NHM) and collected (USNPC) and fit a break-point regression using the `segmented` package for R. (64)

#### Body size

We examined trends in body size of hosts and parasites over time using the date of description given in the NHM dataset. For parasite body size, we used a recently-published database of trait information for acanthocephalans, cestodes, and nematodes (65), and recorded the adult stage body length for all species present in the NHM dataset. For host body size, we subsetted associations to mammals with body mass information in PanTHERIA (66). We examined trends in worm length and host mass over time using generalized additive models (GAMs) with a smoothed fixed effect for year, using the `mgcv` package in R. (67)

#### Host specificity

To test for a description bias in host specificity, we identified the year of description from every species in the NHM data, and coded for each species whether or not they were the first species recorded in the genus. We compared host range for first and non-first taxa and tested for a difference with a Wilcoxon test (chosen given the non-normal distribution of host specificity). To test for temporal trends in host specificity, we fit two GAM models with host specificity regressed against a single smoothed fixed effect for time. In the first, we used the year of species description in the NHM data; in the second, we recorded the year of first accession in the USNPC.

### Estimating Species Richness

Strona and Fattorini (15) discovered that subsampling the host-helminth network produces an approximately power-law scaling pattern, leading to massively reduced richness estimates compared to Dobson *et al*. (1). This pattern was recently found by Carlson *et al*. (14) to be general across large bipartite networks, who developed the R package `codependent` (34) as a tool for fitting these curves and extrapolating symbiont richness. We used the cleaned host-helminth network and `codependent` to fit curves for each of twenty groups, and extrapolate to independent richness estimates for all host groups. We sourced the estimate of every terrestrial group’s diversity from the 2014 IUCN Red List estimates. Fish were split into bony and cartilaginous fish in the same style as Dobson *et al*. (1), but because they have much poorer consolidated species lists, we used estimates of known richness from a fish biology textbook. (68)

The software also allows generation of 95% confidence intervals generated procedurally from the fitting of the networks, and while we have used these in previous work (14), here we elected not to. In our assessment, the epistemic uncertainty around cryptic species, the percent of documented links, and even basic choices like the number of bony fish far outweigh the uncertainty of the model fit for the power law curves.

One major methodological difference between Carlson *et al*. (14) and our study is that in their study, they back-corrected estimates by the proportion of viruses described for the hosts in their network (via validation on independent metagenomic datasets). We have no confident way to evaluate how comprehensive the NHM dataset is, as it is certainly the largest dataset available describing host-helminth interactions, and widely believed to be one of the most thorough. (6) Consequently, our estimates account for the proportion of undescribed diversity due only to unsampled hosts, and underestimates by assuming all recorded hosts have no undescribed parasites. This error is likely overcorrected by the back of the envelope correction we perform for cryptic richness.

### Estimating Total Richness Across Host Groups

The overall number of parasites for all orders considered is smaller than the sum of estimates for each order, as some parasites would be expected to infect vertebrates from more than one order. Here we present a new mathematical approach to correcting richness estimates for affiliates across multiple groups, based on the inclusion-exclusion principle.

#### Inclusion-Exclusion Principle

The inclusion-exclusion principle from set theory allows us to count the number of elements in the union of two or more sets, ensuring that each element is counted only once. For two sets, it is expressed as follows:

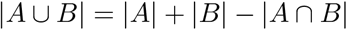

Where |*A* ∪ *B*| is the number of elements in the union of the set, |*A*| and |*B*| are the number of elements in A and B, respectively, and |*A* ∩ *B*| is number of elements in both A and B. For three sets, it is expressed as follows:

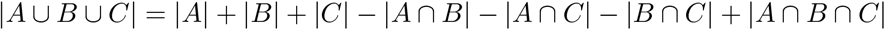

For a greater number of sets, the pattern continues, with elements overlapping an even number of sets subtracted, and elements overlapping an odd number of sets added.

#### Inclusion-Exclusion and Parasite Estimates

The overall estimated number of parasites of two groups, 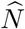, is given as the expected size of 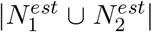. Adapting the inclusion-exclusion principle, we can assume that the overlap between groups *N*_1_ and *N*_2_ in collections is similar to the overlap of not yet discovered parasites:

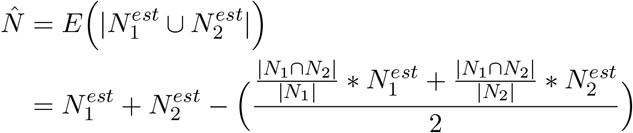

We average the estimated number in both groups over 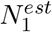 and 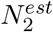, rather than just scaling by |*N*_1_ ∩ *N*_2_|/(*N*_1_ + *N*_2_), because we cannot be sure that 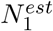 and 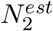 scale with *N*_1_ and *N*_2_ roughly proportionally. (For example, we estimated that the description rate of mammal trematodes is almost an order of magnitude higher than in reptiles.) Instead of estimating the average overlap for a given total number, we estimate the number of multi-order parasites for a given order’s count, and average that across the groups.

For *h* orders, this can be generalized as follows:

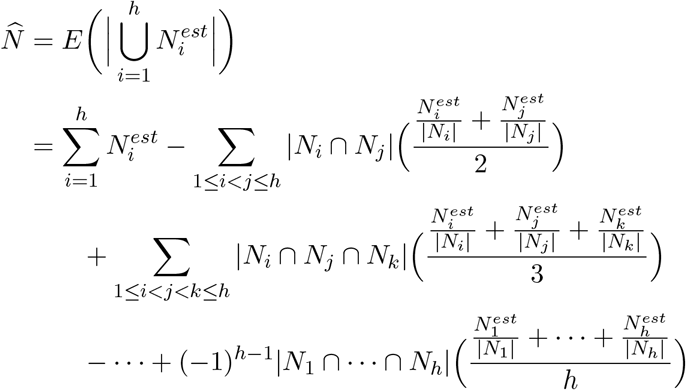

We provide a new implementation of this approach with the `multigroup` function in an update to the `R` package `codependent`.

### Mapping Potential Richness

To map species richness, we used the IUCN range maps for mammals, and counted the number of mammals overlapping each country. Using mammal richness for each country, we predicted the expected number of parasitic associations those species should have globally, running models separately by parasite group (acanthocephalans, cestodes, ne-matodes, and trematodes), and totalled these. We call these “possible” associations and not expected richness, for two reasons: (1) Most macroparasites, especially helminths, are not found everywhere their hosts are found. (2) Host specificity may vary globally (69), but as we stress in the main text, it is difficult to disentangle our knowledge of macroecological patterns from the massive undersampling of parasites in most countries. We compared patterns of possible richness against known helminth associations recorded in a given country, the grounds on which parasite richness has previously been mapped. (6) Finally, we mapped the percentage of total possible unrecorded interactions (an upper bound for high values, except when 100% is reported, indicating that no parasites have been recorded in the NHM data from a country).

